# LATE MERISTEM IDENTITY2 regulates cuticle formation on the seed surface and influences seed longevity

**DOI:** 10.1101/2024.11.14.623493

**Authors:** Yoshimi Oshima, Takako Narumi, Yasuko Kaneko, Toshiki Ishikawa, Maki Kawai-Yamada, Masaru Ohme-Takagi, Nobutaka Mitsuda

## Abstract

The seed coat protects the embryo from severe external environment. Like the aerial plant parts, the exterior surface of the seed coat is surrounded by a cuticle consisting of lipid polyesters and wax. However, the protective role of the cuticle in mature seeds has not been characterized. We herein study the transcriptional regulation of cuticle formation on the seed surface, which influences seed longevity. A chimeric repressor of *Arabidopsis thaliana* R2R3 MYB transcription factor gene *LATE MERISTEM IDENTITY2* (*LMI2)* and a *Torenia fournieri* ortholog (*TfMYBML3*) was expressed in *A. thaliana* and *T. fournieri*, respectively. These lines and *A. thaliana LMI2* T-DNA insertion lines were analyzed for cuticle deficiencies. *A. thaliana LMI2* mutants and chimeric repressor lines exhibited a similar cuticle deficiency in immature seeds. Transmission electron microscopy observations, polyester composition analyses, and gene expression investigations revealed that *LMI2* plays an essential role in cuticle development in the seed coat epidermis by regulating cutin biosynthesis. The *lmi2-2* mutant exhibited shorter seed longevity than the wild-type controls in an accelerated aging test. These results indicate that LMI2 is a key regulator of cuticle formation in the seed coat, and is important for maintaining seed longevity.

**Significance statement:** Cuticle is well known to have multiple important roles for surviving in the terrestrial environment, but their specific role on the seed coat and transcriptional regulation have not yet been elucidated. Here we show that *LATE MERISTEM IDENTITY 2*, encoding a transcription factor regulating floral meristem transition, is required for seed cuticle formation and maintaining seed longevity, providing evidence for a link between surface cuticle and seed longevity.

## Introduction

Orthodox seeds survive over long periods at low temperatures and humidity although seed viability and vigor deteriorate if they are exposed to biotic stresses and/or experience metabolic changes, such as oxidation, or cellular damage (Desai, 2004; Rajjou & Debeaujon, 2008). Because 90% of crops propagate by seeds, maintaining seed viability is agronomically and economically important (Desai, 2004).

In part because of its impermeability and pigments, the seed coat (testa) protects the embryo from pathogens and predators, UV radiation, moisture, high temperatures, and other environmental stresses (Rajjou & Debeaujon, 2008). The *Arabidopsis thaliana* seed coat develops from maternal tissues consisting of the chalazal region and the outer and inner integuments of the ovule. It differentiates into five cell layers, which contain different compounds. The innermost layer (endothelium) accumulates a proanthocyanidin called tannin, which produces a brown mature seed, and a lipidic polyester layer, cutin (Haughn and Chaudhury, 2005; Molina *et al*., 2008; Panikashvili *et al*., 2009). Another lipid polyester, suberin, accumulates in the cell wall of outer integument 1 (Molina *et al*., 2008; Franke *et al*., 2009). The seed coat epidermis differentiates into special cells containing volcano-shaped secondary cell walls called columella and pectinaceous mucilage, which promotes seed germination by retaining water. In the later stages of seed development, all seed coat cells die and are combined (Haughn & Chaudhury 2005; Western *et al.,* 2000). Among the major seed coat components, suberin and tannin help make the seed coat impermeable to water (Debeaujon *et al*., 2000; Beisson *et al*., 2007; Molina *et al*., 2008).

Cutin (lipid polyester) also accumulates on the seed coat surface together with wax to produce the cuticle, which prevents organ fusions between ovules and seeds (Panikashvili *et al*., 2009; Panikashvili *et al*., 2010; Tanaka *et al*., 2007; Watanabe *et al*., 2004). Mutations to a BAHD acyltransferase, DEFECTIVE IN CUTICULAR RIDGES (DCR), which has shown in vitro diacylglycerol acyltransferase activity and localization in the cytosol, lead to fused seeds as well as petal and leaf cuticle deficiencies (Panikashvili *et al*., 2009). However, it is unclear whether the cuticle has a protective role in mature seeds.

We previously reported that the MIXTA-like transcription factors (TFs), MYB106 and MYB16, synchronously regulate epidermal cell morphogenesis and cuticle development (Oshima *et al*., 2013a, Oshima *et al*., 2013b). Single and double knockout/downs of *MYB106* and *MYB16* result in defective (e.g., reduced nanoridges on petals and filaments) and permeable cuticles, suggesting the overlapping roles of MYB106 and MYB16 in cuticle formation (Oshima *et al*., 2013a, Oshima *et al*., 2013b). However, the seed phenotype of these mutants was relatively normal. Therefore, it is possible that other factors regulate cuticle development in seeds.

In this study, we focused on the LATE MERISTEM IDENTITY2 (LMI2) TF. The MYB106/MYB16 TFs and LMI2 belong to neighboring clades (Oshima *et al*., 2013a). Although the proteins of both clades commonly include an R2R3-type MYB domain and an adjacent domain characteristic of subgroup 9 (Stracke *et al*., 2001), there are apparent differences in the C-terminal domain structure. The LMI2 TF induces the floral identity pathway through the direct activation of APETALA1 (AP1), and represses the inflorescence identity pathway by forming a heterodimer with LEAFY (LFY) (Pastore *et al*., 2011). Knockout plants lacking *LMI2*, namely *lmi2-1* and *lmi2-2*, exhibit increased production of secondary inflorescences, and an enhanced *lfy-10* phenotype, including delayed transition of meristem identity (Pastore *et al*., 2011). These findings suggest that the two clades of the R2R3 MYB family subgroup 9 are functionally different. However, in this study, the *LMI2* T-DNA insertion line exhibited defective cuticle development on the seed coat epidermis, which decreased seed longevity and caused seeds to fuse. Our study demonstrates that the seed cuticle is an important regulator of seed longevity, and that the dual-functioning LMI2 TF regulates floral meristem transition and cuticle formation on the seed surface.

## Results

### Constitutive expression of the *LMI2* chimeric repressor resulted in cuticle deficiencies

We first constructed a *LMI2* chimeric repressor, in which *LMI2* is fused to the *SRDX* repression domain (*LMI2-SRDX*) under the control of the constitutive promoter *CaMV 35S*. In plants, chimeric repressors suppress the expression of the target genes of each TF, resulting in loss-of-function phenotypes related to the TFs and any redundant TFs (Hiratsu *et al*., 2003). In our phenotypic analysis of the chimeric repressor line (*35S:LMI2-SRDX*), we observed increase of secondary florescence similar to a T-DNA tagged line of *LMI2* mutant (*lmi2-2,* Figure 2(a)), suggesting that the chimeric repressor line reproduced loss of function of *LMI2* (Figure S1). Additionally, we found fusions between leaves and between floral organs, which were stacked in buds (Figure S2(a–f)). Among 21 T_1_ transgenic plants, 12 exhibited varying degrees of organ fusion (Table S1). The organ fusion phenotypes were almost identical to those of the *MYB106* chimeric repressor lines, which produced defective cuticles (Oshima *et al*., 2013a). In the mild- phenotype lines, there was a reduction in the abundance of epicuticular wax crystals on the inflorescence stem (Figure S2(g,h)). We also examined the water permeability of the leaf surface using a toluidine blue (TB) test, in which plant tissues with a defective cuticle were stained with the water soluble TB dye that cannot penetrate intact cuticles (Tanaka *et al*., 2004). The *35S:LMI2-SRDX* seedlings were stained blue and contained significantly higher amounts of TB than the wild-type seedlings, which were not stained by TB (Figure S2(i–k)).

LMI2 belongs to a different clade from the one including MYB106 and MYB16 regulating cuticle development in MYB subgroup 9 and is known to be involved in floral meristem identity (Oshima *et al*., 2013a; Pastore *et al*., 2011). Indeed, over-branched trichomes observed in mutants and chimeric repressor lines of MYB106 (Oshima *et al*., 2013a) were not found in *35S:LMI2-SRDX* plant and *lmi2-2* although less-branched trichomes were slightly increased in the *35S:LMI2-SRDX* and *lmi2-2* plants, suggesting that protein function of LMI2 is different from MIXTA-like TFs (Table S2). We cloned a putative ortholog of *LMI2*, *TfMYBML3*, from *Torenia fournieri* (Figure S3) and investigated whether it also functions in cuticle development. The leaves of transgenic *T. fournieri* plants expressing *35S:TfMYBML3-SRDX* were clearly stained by TB (Figure S2(l,m)), suggesting that the cuticle development-related functions of LMI2-clade MYBs are not specific to *A. thaliana*.

### *LMI2* promoter activity in immature seed coats and silique tissues

*LMI2* is expressed throughout the shoot apical meristem of primary inflorescences and the carpels and stamens of the subsequent floral development stage. Additionally, the floral organs do not fuse in *lmi2* mutants (Pastore *et al*., 2011). *LMI2* is also highly expressed in siliques (Zhang *et al*., 2009). To reveal the tissue-specific *LMI2* expression patterns during the early silique developmental stages, we conducted promoter–reporter analyses using β-glucuronidase (GUS) and green fluorescent protein (GFP) as reporters. A highly active *LMI2* promoter was observed in immature seeds, septa, and funicles, as well as in the dehiscence zone (Figure 1(a–c)). Confocal laser scanning microscopy observation revealed that the *LMI2* promoter was clearly active in outer integuments from 0 to 7 day after flowering (DAF) and in inner integument 1 layer from 1 to 5 DAF (Figure 1(d)). Other inner integument layers also had weak activity of *LMI2* promoter from 0 to 2 DAF (Figure 1(d)). *LMI2* promoter used in reporter experiments was sufficient to complement the *lmi2-2* mutation when *LMI2pro:LMI2-GFP:HSPt* was transformed as described later (Figure 2). These results suggest that LMI2 functions in the epidermis of immature seed coats and silique tissues during early seed developmental stages.

**Figure 1.**
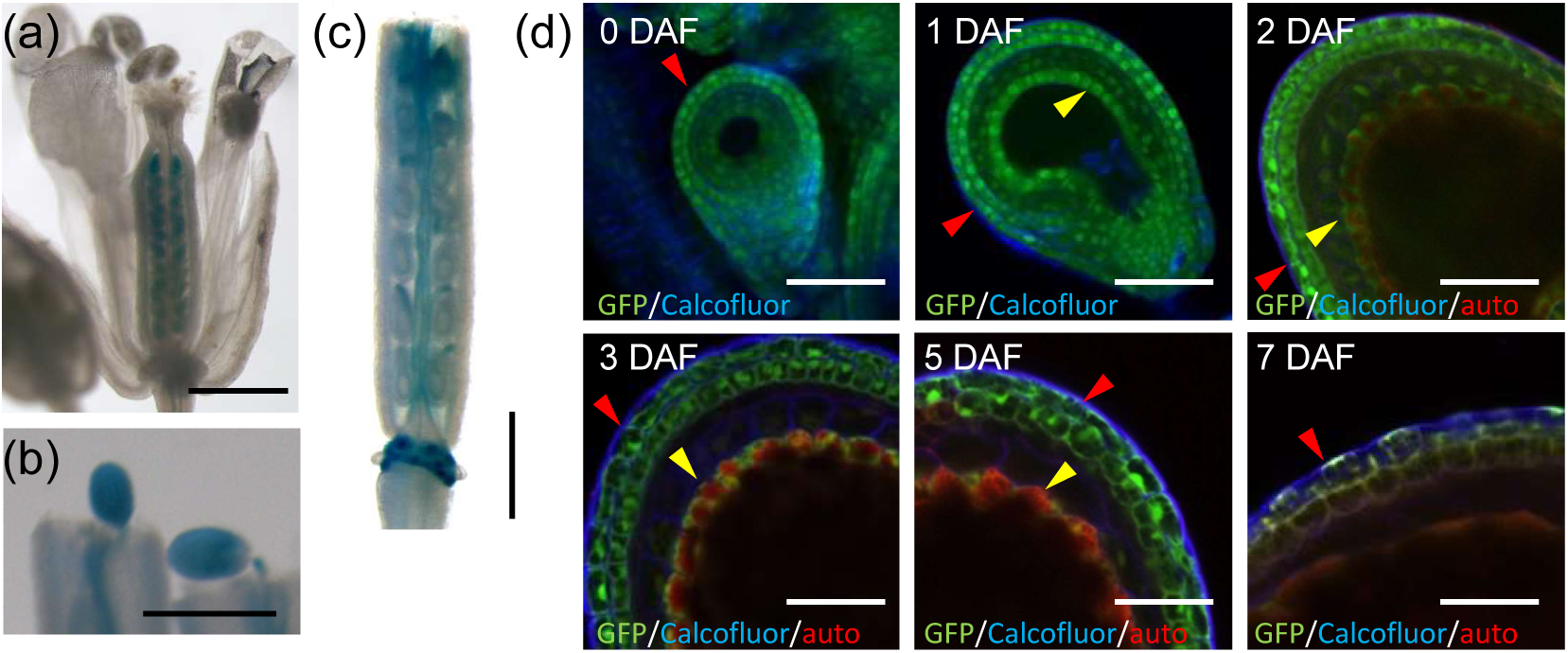
*LMI2* promoter activity. (a, b and c) β-glucuronidase reporter activity (blue stain) driven by the *LMI2* promoter in a flower (a), immature seed (b), and immature silique (c). (d) Green fluorescent protein (GFP) reporter activity driven by the *LMI2* promoter in a seed on the day the flower opened (0 DAF), 1, 2, 3, 5 and 7 day after flowering (1, 2, 3, 5 and 7 DAF). GFP, calcofluor white and auto fluorescence images are merged. Red and yellow arrow heads indicate GFP localization in seed coat outer integument 2 (epidermis) and inner integument 1 (endothelium), respectively. Bars indicate 500 μm in (a), (b), and (c), and 50 μm in (d).

**Figure 2.**
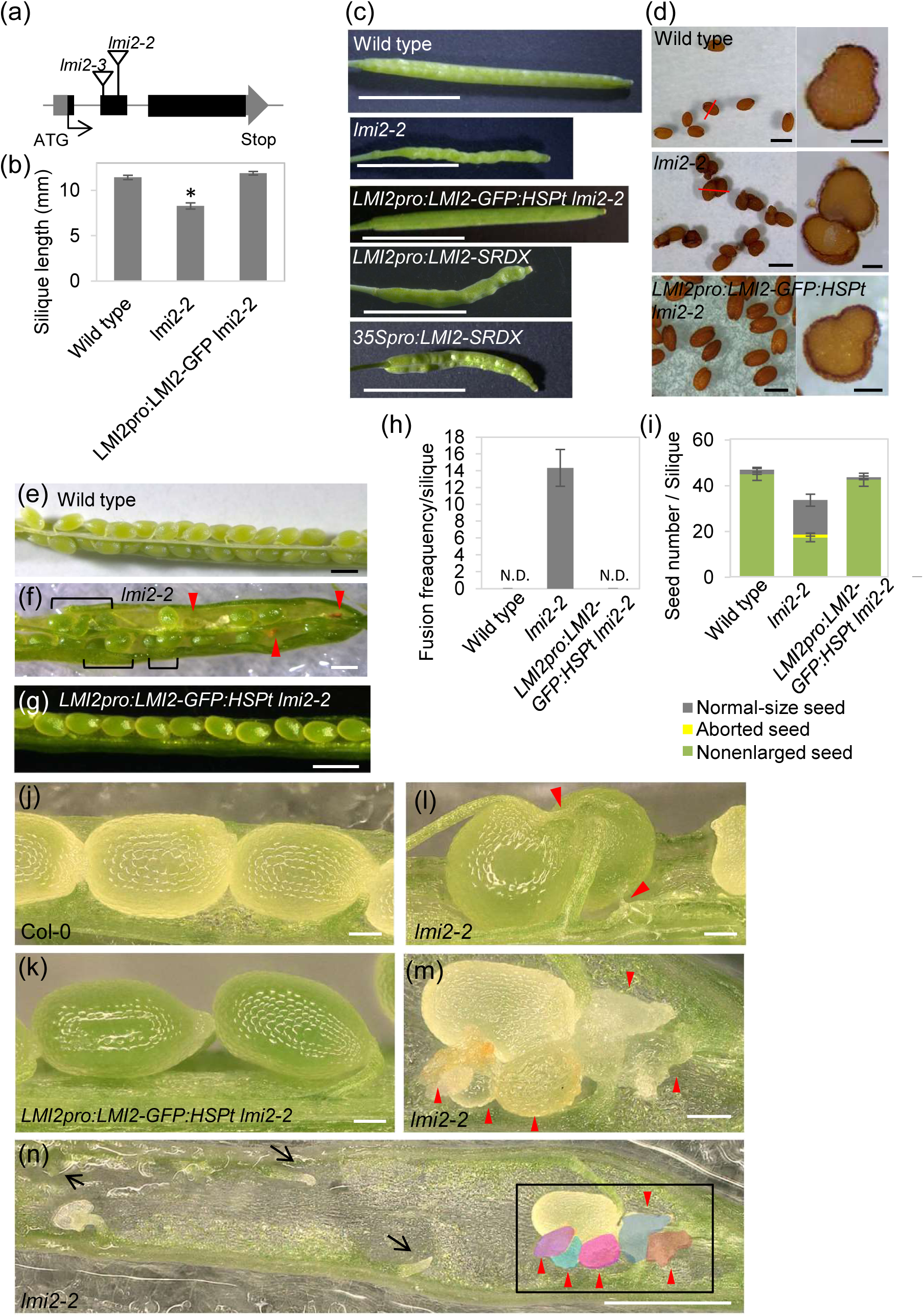
Phenotypes of the *lmi2-2* mutant. (a) Structure of the *LMI2* gene and the position of the T-DNA insertions. (b) Average length of mature siliques on the main stem of wild-type and *lmi2-2* plants. Error bars represent standard errors (n = 82 or 97). Asterisks represent *P* < 0.01 according to Welch’s *t*-test. (c) Representative wild-type, *lmi2-2*, *LMI2pro:LMI2-GFP:HSPt lmi2-2, LMI2pro:LMI2-SRDX* and *35Spro:LMI2-SRDX* silique with average lengths. (d) Mature seeds of wild-type, *lmi2-2* and *LMI2pro:LMI2-GFP:HSPt lmi2-2* plants. Cross sections of the sites indicated by a red line in the left panels are presented. Fusion of *lmi2-2* seeds occurred outside of the seed coat. (e, f, g, j to n) Internal contents of wild-type, *lmi2-2* and *LMI2pro:LMI2-GFP:HSPt lmi2-2* immature siliques. Red arrowheads and square brackets indicate fused seeds. Solid-line arrows indicate torn funicles. The region in a square (n) is magnified in (m). (h) Number of fusions in a wild-type, *lmi2-2,* and *LMI2pro:LMI2-GFP:HSPt lmi2-2* silique. Error bars represent standard errors (n = 9). Asterisks represent *P* < 0.01 according to Welch’s *t*-test. N.D. indicates not detected. (i) Number of normal-size seeds, aborted and not-enlarged seeds in one wild-type, *lmi2-2* and *LMI2pro-LMI2-GFP:HSP lmi2-2* immature silique. Error bars represent standard errors (n = 9). The number of normal (*P* < 0.01) and not-enlarged (*P* < 0.05) size seeds are significantly different between wild-type and *lmi2-2* plants according to Welch’s *t*-test. Bars indicate 5 mm in (c), 500 μm in (e), (f), (g), (n) and left panel of (d), and 100 μm in (j) to (m) and right panel of (d).

### Loss of *LMI2* alters seed coat cuticle formation

To investigate the biological functions of LMI2 in immature siliques and seeds, we analyzed the chimeric repressor lines expressed under the original promoter, *LMI2pro:LMI2-SRDX*, and two independent lines carrying *LMI2* disrupted by a T-DNA insert (i.e., *lmi2-2* and *lmi2-3*; Figure 2(a)) (Pastore *et al*., 2011). The *LMI2pro:LMI2- SRDX*, *lmi2-2* and *lmi2-3* plants consisted of slightly shortened siliques with a rough surface (Figure 2(b,c), Figure S4(a,d)). *35S:LMI2-SRDX* plants also showed similar phenotypes (Figure 2(c)). We also observed fusions between seeds as well as decreased seed number in siliques (Figure 2(d–n), Figure S2(o), S4(b,e,f)). The seeds in immature siliques were clustered, and some of fused seeds were smaller than unfused seeds (Figure 2(e,f,i,l,m), Figure S4(b,c)). Torn or stretched funicles surrounding fused seeds were common, implying that a lack of functional funicles to transport nutrients led to seed growth defect (Figure 2(n), Figure S4(c)). All these phenotypes of *lmi2-2* was complemented by *LMI2* expression under the *LMI2* promoter (Figure 2(b, c,g,-i,k)). Because *LMI2pro:LMI2-SRDX*, *lmi2-2* and *lmi2-3* plants produced similar phenotypes, we subsequently focused on the *lmi2-2* line.

To investigate whether maternal surface defect caused seed reduction, we crossed *lmi2-2* and wild-type plants. When *lmi2-2* carpel was pollinated with wild-type pollens, the F_0_ siliques and F_1_ seeds showed rough surface short phenotype and fusion, respectively. However, when wild-type carpel was pollinated with *lmi2-2* pollens, the F_0_ silique and F_1_seed phenotypes were similar to wild type (Figure 3(a), Figure S5(a,c,d,f)). Both F_1_ plants were heterologous genotype and produced normal siliques and seeds (Figure 3(a), Figure S5(b,c,e,f,g)). Seed number is reduced if the maternal genotype was *lmi2-2* (Figure 3(b,c)). Even though the quarter of F_2_ seeds in F_1_ silique are *lmi2-2* homo, heterologous F_1_ plants produced similar number of seed to wild type (Figure 3(c)). These results suggest that seed reduction is caused by maternal tissue defect including surface fusion between seed coat, funicle and silique tissue.

**Figure 3.**
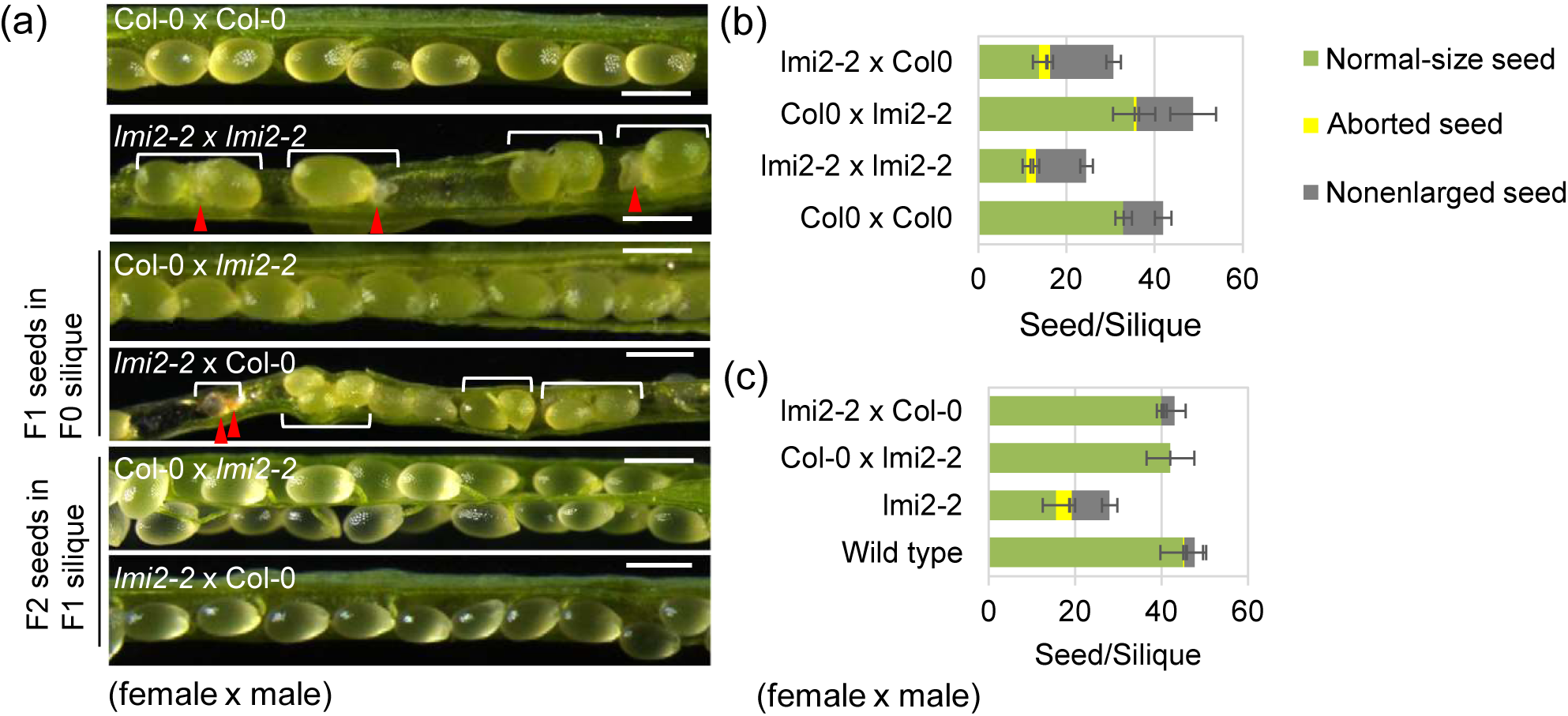
Phenotypes of immature seeds in the F_1_ and F_2_ generation of wild type and *lmi2-2* mutant. (a) Internal contents of siliques in F_0_ and F_1_ plants of indicated “female (left) x male (right)”. Red arrowheads and square brackets indicate fused seeds. (b and c) The number of normal, aborted and not-enlarged seeds in one immature silique in F_0_ (b) and F_1_ (c) plants of indicated combinations. Error bars represent standard errors [n = 10 for (b), n = 3 for (c)]. The number of normal and aborted seeds is significantly different between Col.0 x Col.0 and *lmi2-2* x *lmi2-2* and between Col.0 x Col.0 and *lmi2-2* x Col.0 (*P* < 0.05 according to Welch’s *t*-test). Bars indicate 5 mm in (a).

The mucilage extrusion pattern of *lmi2-2* mature seeds was similar to that of wild-type seeds, except for the immature size seeds (Figure S6(a–c)). The failure of the seed coat cell layer to develop, loss of epidermis, and thickened outer integument one cell layer were also observed in a cross section of the fused region, suggesting that the fusion occurred before cell differentiation (Figure S6(d,e)). The TB test revealed that the surfaces of the septum and immature seed were permeable, indicating a cuticular functional deficiency (Figure 4(a–d)). The structure and thickness of the seed coat cuticle were observed by transmission electron microscopy (TEM) at 1 DAF, which is when *LMI2* is being expressed, and at 8–10 DAF (i.e., bent-cotyledon stage) when fused seeds were detected. At 1 DAF, the cuticle of wild-type seeds consisted of an electron-dense layer outside of the primary cell wall of the seed coat epidermis (Figure 4(e)). During the bent-cotyledon stage in wild-type samples, the primary cell wall was covered with a thick electron-dense cuticle layer (Figure 4(h)). In contrast, *lmi2-2* samples occasionally lacked or had a thin loosely structured electron-dense layer covered by an amorphous layer (Figure 4(f,g,i,j)). On average, the electron-dense layer was significantly thinner in *lmi2- 2* seeds compared with the corresponding layer in wild-type seeds (Figure 4(k,l)). The cuticle of mature seeds is difficult to observe after chemical cross-linking because aqueous solutions induce the extrusion of mucilage and disruption of the cuticle. We observed the cuticle surrounding mature seeds using a scanning electron microscope in backscattered electron (BSE) mode, which gives brightness depending on chemical composition. A brightly colored layer on the seed surface in BSE images was observed (Figure 4(m)). When the same samples were stained with Fluorescent Brightener 28, which stains cellulose, the corresponding layer located on the exterior of cell wall emitted yellow auto fluorescence, indicating that the layer is cuticle (Figure S7). These findings suggest that the brightly colored layer represented the cuticle. This layer was thinner in mature *lmi2-2* seeds than in wild-type seeds (Figure 4(l)). Our observations clearly indicate that cuticle formation in the seed coat epidermis of the *lmi2-2* mutant is incomplete.

**Figure 4.**
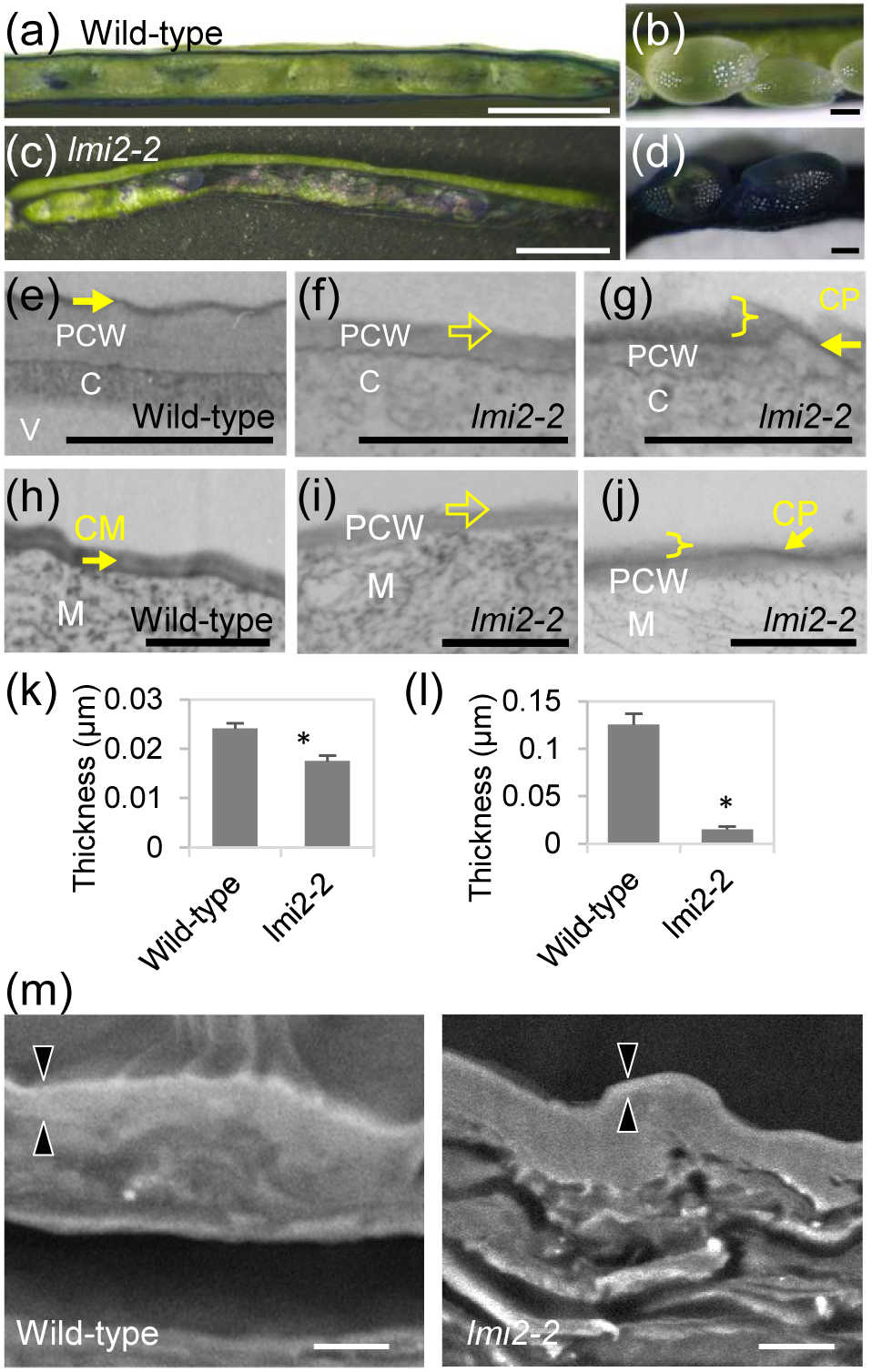
Cuticle formation in *lmi2-2* mutant seeds and siliques. (a and c) Immature silique of wild-type (a) and *lmi2-2* (c) plants from which one side of the valves and seeds were removed and stained with toluidine blue. The site between the replum and valve pierced by a needle was commonly stained blue. The septum was differentially stained in wild-type and *lmi2-2* samples. (b and d) Immature seeds of wild-type (b) and *lmi2-2* (d) plants stained with toluidine blue. (e, f, g, h, i and j) Transmission electron micrograph of a cross section of the seed coat epidermis in wild-type (e and h) and *lmi2-2* (f, g, h and i) at first day after flowering (1 DAF) (e, f and g) and at the bent-cotyledon stage (h, i and j). Filled arrows indicate the electron dense layer of the cuticle proper (CP) or cuticle membrane (CM) as in the wild-type sample. Open arrows indicate a lack of cuticle. A brace indicates the cuticle proper with an electron-lucent amorphous outer layer. PCW, primary cell wall; C, cytosol; V, vacuole. (k and l) Average thickness of the cuticle electron dense layer in wild-type and *lmi2-2* seeds at 1 DAF (k) and at the bent-cotyledon stage (l). Error bars represent standard errors (n = 4 or 5). Asterisks indicate *P* < 0.05 according to Welch’s *t*-test. (m) Scanning reflection electron micrograph of a mature seed coat of wild-type and *lmi2-2* seeds. Pair of black arrowheads indicates the surface cuticle. Bars indicate 1 mm in (a) and (c), 100 μm in (b) and (d), 1 μm in (e), (f), (g), (h), (i) and (j), and 3 μm in (m).

### Cutin biosynthesis is suppressed in the *lmi2* mutant

Mature seed coat cuticle mainly consisted of lipid polyesters and waxes. We extracted the polyesters, including the embryo cuticle, and three types of polyester derived from the seed coat, cuticular cutin, and suberin that accumulated in outer integument layer 2 as well as the cutin that accumulated in the inner integument. Extracted polyesters were depolymerized and analyzed by gas chromatography–mass spectrometry. The composition of polyester monomers differed between the wild-type and *lmi2-2* seeds, although the total abundance of polyester monomers were slightly decreased (Figure 5(a,b)). The C18–22 alcohols and sinapate contents were lower, in *lmi2-2* seeds. Especially, 10, 16-dihydroxy fatty acid in *lmi2-2* was decreased by a quarter of wild type. The 18:1 and 18:2 ω-hydroxy fatty acid contents of the *lmi2-2* seeds were slightly higher and lower than the wild-type levels, respectively. The compositional ratio of C16-24 dicarboxylic acids were comparable to wild type (Figure 5(b)). We also extracted cuticular waxes from the surface of mature seeds for analysis by gas chromatography– mass spectrometry. The total amount of wax monomers was higher in *lmi2-2* seeds than in wild-type seeds (Figure 5(c)). All of monomer contents in *lmi2-2* were differentially increased except for C33 alkane, C16 and C18 fatty acids (Figure 5(c,d), Figure S8). The difference in the abundance of wax monomers may be related to the amorphous layer covering the electron-dense cuticle layer in *lmi2-2* seeds. These results suggest that *LMI2* is involved in polyester biosynthesis, which affects the accumulation of waxes.

**Figure 5.**
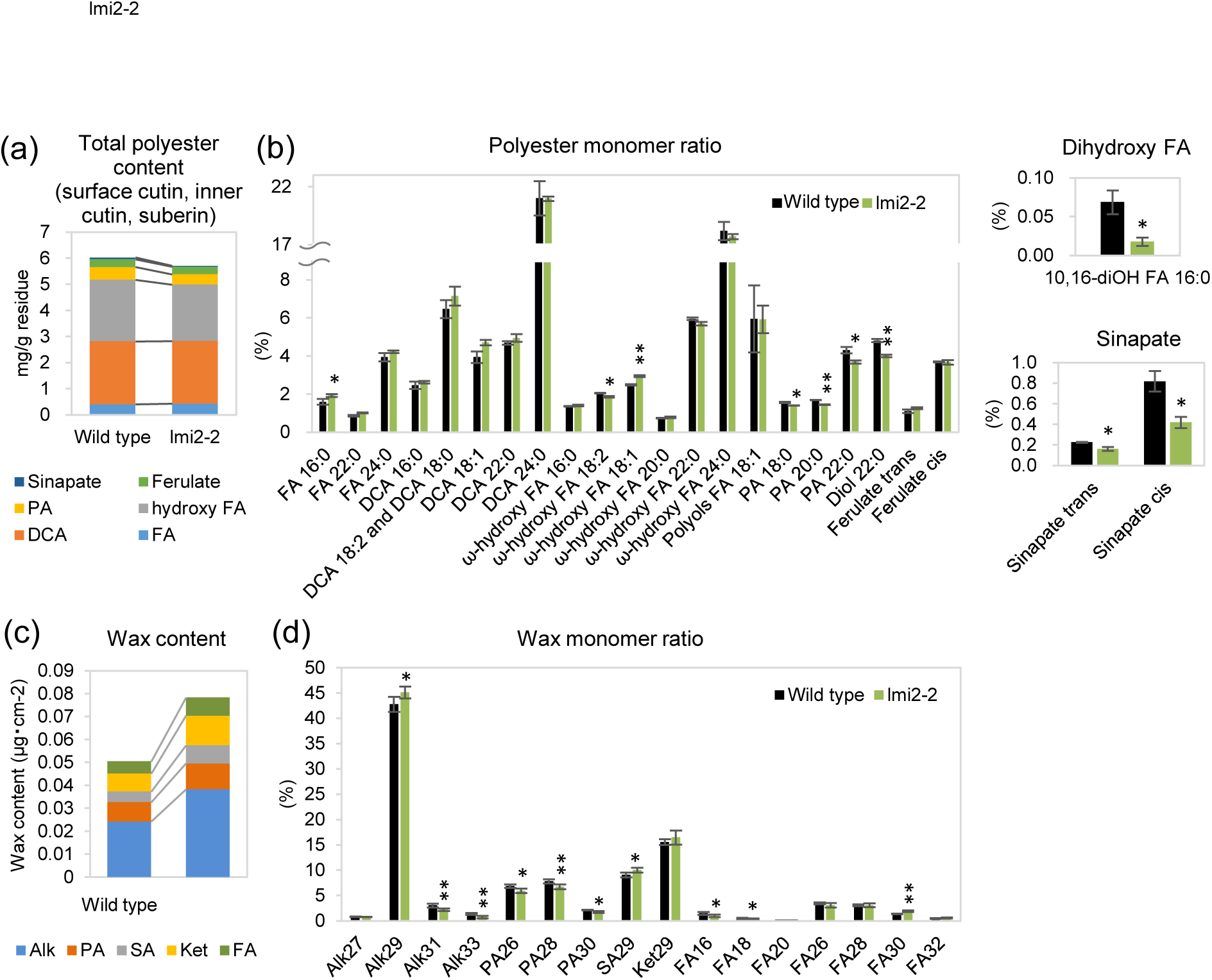
Wax and polyester content of seeds. (a) Wax content in wild-type and *lmi2-2* mature seeds. Alk; alkanes, PA; primary alcohols, SA, secondary alcohols, Ket; ketone, FA; fatty acids (b) Wax monomer ratio in wild-type and *lmi2-2* mature seeds. Error bars represent standard error (n = 6 or 5). (c) Total polyester content in wild-type and *lmi2-2* mature seeds. Hydroxy FA; hydroxy fatty acids, DCA; dicarbonic acids (d) Polyester monomer ratio in wild-type and *lmi2-2* mature seeds. Error bars represent standard error (n = 3). Single and double asterisks represent *p*<0.05 *p*<0.01 by Welch’s t-test, respectively.

To investigate how polyester biosynthesis is regulated by *LMI2*, we analyzed the expression of genes differentially expressed in the seed coat and during developmental stages. We extracted total RNA from siliques at 1 DAF and from immature seeds at 7 DAF, and analyzed gene expression levels by quantitative reverse transcription polymerase chain reaction (qRT-PCR). The expression levels of *DCR* and *CYP77A6*, which are involved in cutin biosynthesis, were lower in *lmi2-2* seeds than in wild-type seeds at 1 DAF. The expression level of the cutin transporter gene, *ABCG13*, was also lower in *lmi2-2* seeds. At 7 DAF, the *lmi2-2 CYP77A6* and *ABCG13* expression levels were not significantly different from the wild-type levels, but the *DCR* expression level in *lmi2-2* seeds was lower than that of the wild-type seeds (Figure 6). The expression levels of *ABERRANT INDUCTION OF TYPE THREE GENES* (*ATT1*) and *GPAT5*, which influence the accumulation of cutin in the inner endothelium and suberin in outer integument 1, respectively, were not suppressed in *lmi2-2* seeds (Figure 6). The expression of *MUM4*, which is required for mucilage biosynthesis and cytoplasmic rearrangement (Western *et al*., 2001; Western *et al*., 2004), was also similar between *lmi2-2* and wild-type samples (Figure 6). These results suggest that LMI2 is involved in cutin biosynthesis in the outer integument during the early seed developmental stage, and that the biosynthesis of cutin, suberin, and mucilage in the inner seed parts was not suppressed by a lack of *LMI2*. Overall, the compositional changes in *lmi2-2* seed polyesters seemed to be caused by suppressed cutin biosynthesis related to the outer cuticle, which was compensated by an increase in cuticular wax accumulation.

**Figure 6.**
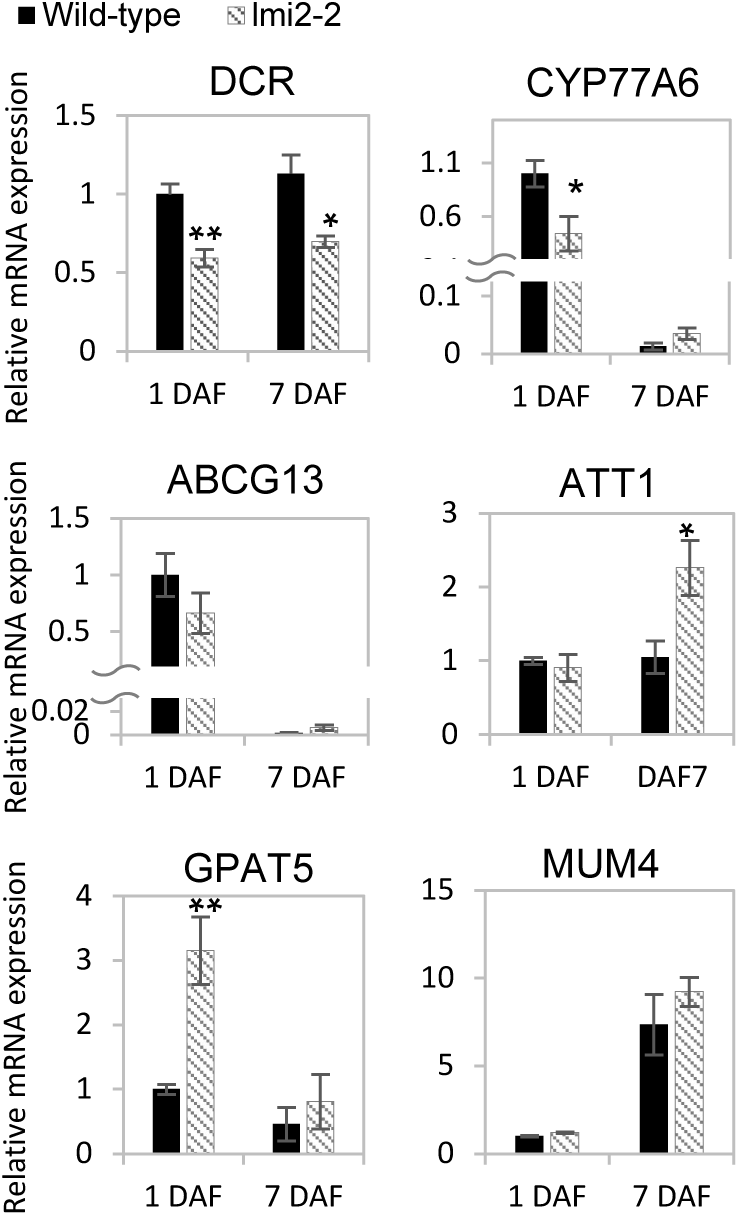
Expression of genes related to the *lmi2-2* seed coat. Quantitative reverse transcription polymerase chain reaction analysis of gene expression in the seed coats of wild-type and *lmi2-2* siliques 1 day after flowering (1 DAF) and in immature wild-type and *lmi2-2* seeds 7 days after flowering (7 DAF). The wild-type expression level is set as 1. Error bars represent standard errors (n = 4). Single and double asterisks indicate *P* < 0.05 and *P* < 0.01 according to Welch’s *t*-test, respectively.

### LMI2 is important for seed longevity

Cutin and suberin in the inner and outer integuments, respectively, were reported to restrict water movement (Molina *et al*., 2008, Panikashvili *et al*., 2009). Mutant seeds with abnormal pigmentation and cell structures have more permeable seed coats, and exhibit decreased longevity (Debeaujon *et al*., 2000). We investigated the biological functions of LMI2 in terms of seed longevity by examining cuticle formation. We conducted an accelerated aging test, in which seeds were subjected to high temperature and humidity for short periods to evaluate relative storability (Delouche & Baskin, 1973). The germination rate of wild-type seeds subjected to accelerated aging conditions was about 20% lower than that of untreated control seeds (Figure 7). In contrast, the germination rate of *lmi2-2* seeds subjected to accelerated aging treatments was approximately 50% lower than that of wild-type seeds (Figure 7). These findings suggest that LMI2 positively regulates seed longevity.

**Figure 7.**
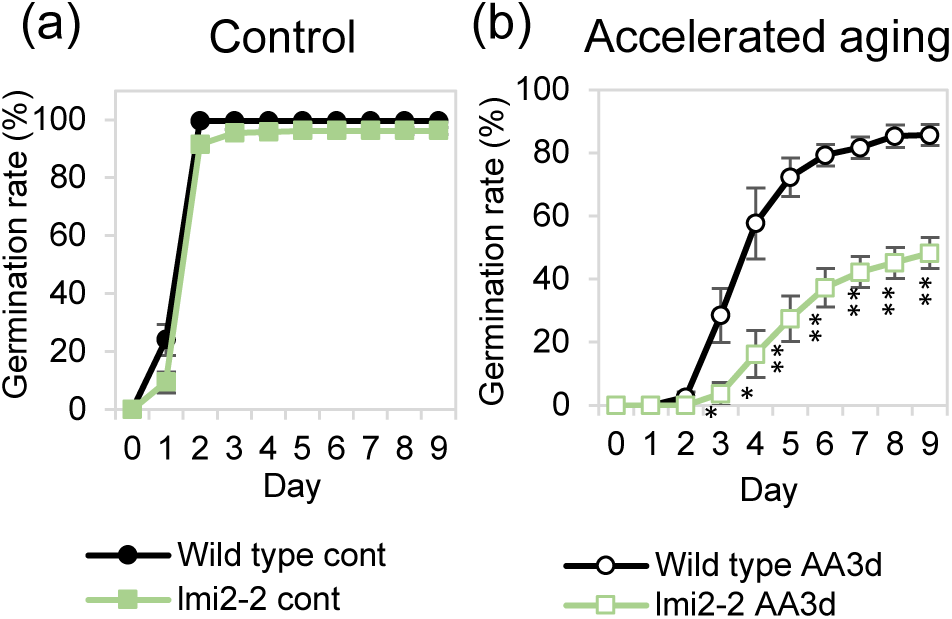
Germination of *lmi2-2* mutant seeds after accelerated aging. Germination rate of wild-type and *lmi2-2* untreated seeds (control) and seeds treated at 40 °C and 100% relative humidity for 3 days (accelerated aging). Error bars represent standard errors (n = 4). Single and double asterisks indicate *P* < 0.05 and *P* < 0.01 according to Welch’s *t*-test, respectively.

To investigate whether maternal surface defect caused seed deterioration, we performed an accelerated aging test for the F_2_ seeds obtained by cross between *lmi2-2* and wild-type plants. The germination rate of F_2_ seeds subjected to accelerated aging conditions was similar to that of wild-type seeds, even though the F_2_ seeds include *lmi2-2* homo (Figure S9). This result suggests that the defect of seed longevity in *lmi2-2* is associated with maternal genotype.

## Discussion

### LMI2 regulates the accumulation of cuticular substances

Cuticle development is regulated by different plant developmental stages and exposure to various stresses. In this study, we revealed a previously unknown LMI2 function related to the regulation of cuticle formation. Constitutive expression of *LMI2- SRDX* led to cuticle deficiencies characterized by organ fusion, decreased epicuticular wax crystal abundance, and increased cuticle permeability essentially throughout the plant. In contrast, mutation in *LMI2* did not decrease wax abundance, but suppressed cutin biosynthesis, resulting in functional and structural deficiencies of the seed coat surface cuticle. In single mutants, the expression of genes regulated by TFs with the same function as LMI2 would not be suppressed. However, these genes would be suppressed in chimeric repressor lines. Therefore, wax biosynthesis genes may be regulated by LMI2 and other TFs. Based on our results, we propose that LMI2 affects cutin biosynthesis more than wax biosynthesis.

The abundance of total wax was higher in the *lmi2-2* mutant than in the wild- type seeds. The over-accumulation of wax has also been reported for the *bdg* mutant, which has a mutation in the gene encoding an α/β hydrolase family protein believed to help polymerize carboxylic esters in cuticles (Kurdyukov *et al*., 2006). The cuticle of *bdg* leaves have a thin or irregular shaped electron-dense layer, which may exist as multiple layers or form the middle layer of the cell wall (Kurdyukov *et al*., 2006). Consequently, the increased wax accumulation in the *bdg* mutant may compensate for the loss of cell wall and cuticle integrity (Kurdyukov *et al*., 2006). Additionally, a similar compensatory over-accumulation of wax has been observed in *fiddlehead* and *lacerate* mutants, which exhibit defective biosynthesis of cuticular polyesters, resulting in the production of chemically and/or structurally altered cuticles (Voisin *et al*., 2009). These observations suggest that increased wax accumulation in *lmi2-2* mutants may be due to altered composition of seed polyesters.

### Dual roles of LMI2 in the epidermis

Previous reports indicated LMI2 is a direct target of and interacts with LFY, and the resulting heterodimer activates *AP1* (Pastore *et al*., 2011, William *et al*., 2004). AP1, which is a MADS box TF, and LFY act as key regulators of the transition from an inflorescence meristem to a floral meristem (Huala & Sussex, 1992; Weigel *et al*., 1992). Session *et al*. (2000) reported that *lfy* mutants are fully rescued by the expression of *LFY* in the L1 layer forming epidermis, which induces the expression of homeotic genes, including the class B and C MADS box genes, *APETALA3* (*AP3*) and *AGAMOUS* (*AG*), in the L1 and inner layers. Expression of the *AG*, *AP3*, *PISTILLATA* (*PI*), and *SEPALLATA3* (*SEP3*) MADS box genes under the control of the *ML1* promoter, which is only active in the L1 layer, is sufficient to induce homeotic changes similar to those promoted by constitutive gene expression (Urbanus *et al*., 2010). In general, changes to the L1 layer affect floral organ development. Double mutants involving HD-ZIP IV TF genes (i.e., *PDF2*, *HDG1*, *HDG2*, *HDG5*, and *HDG12*) that mediate the establishment of the L1 layer, produce abnormal flowers with sepaloid petals and carpelloid stamens because of decreased *AP3* expression levels (Kamata *et al*., 2013). Furthermore, the HD-ZIP IV family of TFs regulates protoderm- and epidermis-specific expression of cuticle-related genes in various organs (Abe *et al*., 2001; Abe *et al*., 2003). These observations suggest that transcriptional regulation in the epidermis is closely linked to floral organ development. It is possible that LMI2 affects the formation of the epidermis rather than regulating meristem identity and cuticle development. Identifying additional interacting protein partners for LMI2 may help characterize the functional diversity of LMI2.

### Effects of the seed surface cuticle on seed longevity

The effects of aging on seeds are mitigated by scavenging of reactive oxygen species, damage repair, dormancy, and seed coat impermeability to external factors. Flavonoids, lignin, mucilage, suberin, and cutin in the seed coat influence seed longevity (Rajjou & Debeaujon, 2008). Gene expression analysis revealed that *LMI2* does not regulate the expression of suberin and (inner) cutin biosynthesis genes. Furthermore, seed coat color, and mucilage deposition were unaffected in *lmi2-2* seeds, implying the effects of proanthocyanidin, and mucilage on seed longevity are not influenced by LMI2. These results suggest that LMI2 mediates seed longevity by regulating cuticle accumulation. The current study has uncovered new cuticular functions in the seed coat. Additionally, the endosperm cuticle was reported to be required for normal seed longevity (De Giorgi *et al*., 2015). Further investigation on the cuticle may enable the development of novel strategies to improve long-term seed storage. Enhanced seed storage techniques may be valuable for species conservation as well as the breeding and propagation of high quality crops.

## Experimental procedures

### Plant Materials and Growth Conditions

*Arabidopsis thaliana* ecotype Columbia-0 and *T. fournieri* ‘Crown Violet’ (Aida *et al*., 2000) were used in this study. For the *A. thaliana lmi2-2* and *lmi2-3* mutants, the T-DNA insertion lines SALK_020792 and SALK_115924 were used, respectively. The *A. thaliana* plants were grown at 22 °C in a 16-h light/8-h dark photoperiod. Seedlings were grown on solid Murashige and Skoog medium, and transferred to soil approximately 3 weeks after germination. The *T. fournieri* growth conditions were as previously described (Narumi *et al*., 2011).

### Plasmid construction and Plant Transformation

To construct *35S:LMI2-SRDX* and *35S:TfMYBML3-SRDX*, the protein-coding region of LMI2 and *Torenia fournieri* Lind. *TfMYB-MIXTA-like3* (*TfMYBML3*, GenBank accession number; AB649196) without stop codon was amplified by PCR using the appropriate primers (see Table S3) and was cloned into the *Sma*I site of p35SSRDXG (Mitsuda *et al*., 2006). The protein-coding region of *LMI2* was also cloned into the SmaI site of pLMI2pro_SRDX_NOS_Entry vector to construct *LMI2pro:LMI2-SRDX*. 5’ upstream region (∼3000bp) of *LMI2* amplified by PCR using appropriate primers (Table S3) was cloned into *Asc*I and *Bam*HI site of pSRDX_NOS_Entry vector (Mitsuda et al. 2007) and pGFP_HSP_Entry vector to generate pLMI2pro_SRDX_NOS_Entry vector and *LMI2pro:GFP:HSP* construct, respectively. *LMI2* promoter fused with *LMI2* coding region which was amplified by PCR using appropriate primers (Table S3) from the *LMI2pro:LMI2-SRDX* construct was cloned into *Asc*I and *Sma*I site of pGFP_HSP_Entry vector to generate *LMI2pro:LMI2-GFP:HSP* construct. pGFP_HSP_Entry was generated from pNOS_Entry vector (Sakamoto *et al*., 2016) by substituting nopaline synthase terminator with the heat shock protein 18.2 (HSP) terminator (Nagaya *et al*., 2010). The transgene cassette of the resulting plasmid was transferred into the T-DNA destination vector pBCKH or pBCKK by Gateway LR reaction (Thermo Fisher Scentific, Waltham, MA) (Mitsuda *et al*., 2006). 5’ upstream regions of LMI2 (∼3000bp) was amplified by PCR and cloned into pDONRG-P4P1R (Oshima *et al*., 2011) by Gateway BP reaction (Thermo Fisher Scientific). For the construction of *LMI2pro:GUS*, the cloned promoter fragment of *LMI2* was transferred into R4L1pDEST_GUS_BCKK (Oshima *et al*., 2013a) by Gateway LR reaction. The constructs prepared as described above were used to transform *A. thaliana* and *T. fournieri* plants using a published method (Oshima *et al*., 2013; Narumi *et al*., 2011).

### Scanning electron microscopy

Fresh flower and stem samples were examined using a VE8000 Real 3D system model scanning electron microscope (Keyence, Osaka, Japan) at an accelerating voltage of 1, 2, and 5 kV. The mature seeds were cut with a razor and the cross sections were observed using a scanning electron microscope (S-3400, Hitachi, Tokyo, Japan) in backscattered electron (BSE) mode at 8 kV.

### Staining of plants

*Arabidopsis thaliana* seedlings grown on Murashige and Skoog medium and leaves from *T. fournieri* plants grown in culture pots were stained with TB following the method described by Tanaka *et al*. (2004). The stained samples with eight biological replicates were washed with water and homogenized in cell lysis buffer (TOYO B-Net Inc.,Tokyo, Japan). The supernatants of the cell lysate were used for the absorbance measurements at 650 nm to determine TB amount and for the measurement of the protein amount to normalize the TB uptake. The GUS activity in flowers and immature siliques with three biological replicates was observed by staining as previously described (Oshima *et al*., 2013a).

### Staining of seeds

Whole immature seeds harboring *LMI2pro:GFP:HSP* were cleared with ClearSee and stained with fluorescent brightener 28 (calcofluor white) as described in the previous study (Kurihara et al. 2015). Stained seeds were observed under a confocal laser scanning microscopy LSM 700 (Carl ZEISS, Oberkochen, Germany) using 405 nm and 555 nm lasers with the laser power set as 2.0% and following emission ranges, short pass of 510 nm, long pass of 580 nm and band pass of 493-581 nm for cell wall (fluorescent brightener 28), auto fluorescence and GFP, respectively. Cross sections of mature seeds were stained with fluorescent brightener 28 to observe cell wall. For staining of mucilage, whole seeds were shaken in water and followed by staining with Ruthenium red according to the previous study (Western *et al*., 2000). Four biological replicates were analyzed. Cross section of seeds embedded in resin as described in the method for Transmission Electron Microscopy (TEM) was stained with 0.05% TB.

### Phylogenetic analysis

The evolutionary history was inferred using the Neighbor-Joining method (Saitou & Nei, 1987) after aligning amino-acid sequences of 10 proteins using MAFFT v6.815b (Katoh & Toh, 2008). The bootstrap consensus tree inferred from 1000 replicates (Felsenstein, 1985) is taken to represent the evolutionary history of the taxa analyzed (Felsenstein, 1985). Branches corresponding to partitions reproduced in less than 50% bootstrap replicates are collapsed. The percentage of replicate trees in which the associated taxa clustered together in the bootstrap test (1000 replicates) are shown next to the branches (Felsenstein, 1985). The tree is drawn to scale, with branch lengths in the same units as those of the evolutionary distances used to infer the phylogenetic tree. The evolutionary distances were computed using the Poisson correction method (Zuckerkandl & Pauling, 1965) and are in the units of the number of amino acid substitutions per site. All positions containing gaps and missing data were eliminated from the dataset (Complete deletion option). There were a total of 175 positions in the final dataset. Phylogenetic analyses were conducted in MEGA4 (Tamura *et al*., 2007).

### Transmission electron microscopy

Details regarding sample preparation and TEM have been described elsewhere (Seki *et al*., 2014). Siliques at 1 DAF and fused seeds at the bent-cotyledon stage were cut with a razor and fixed in a solution consisting of 2% (v/v) glutaraldehyde and 0.05 mM potassium phosphate buffer (pH 7.0). Samples were treated with OsO_4_, and the fixed tissues were then dehydrated and embedded. Ultra-thin sections (90–100 nm thick) were prepared for subsequent microscopic analysis. After staining with uranyl acetate and lead citrate, the sections were observed using an H-7500 transmission electron microscope (Hitachi, Tokyo, Japan). To determine cuticle thickness, the TEM photos were analyzed using AxioVision 4.8 software (Carl Zeiss, Jena, Germany).

### RNA analysis

Total RNA was isolated from 1 DAF carpels and 7 DAF seeds with three biological replicates using a modified CTAB method (Sakamoto *et al*., 2012), and then treated with DNase I (Takara Bio, Kusatsu, Japan). Following first-strand cDNA synthesis using the PrimeScript RT kit (Takara Bio), qRT-PCR analysis was conducted according to a published procedure (Oshima *et al*., 2013a). The qRT-PCR primers are listed in Table S3.

### Wax analysis

Cuticular wax of mature seeds from simultaneously grown plants with five biological replicates was extracted by immersing in chloroform containing tetracosane as internal standard for 10 s. The solvent was evaporated in a steam of nitrogen. Free hydroxyl and carboxyl groups were silylated with N,O-Bis(trimethylsilyl)trifluoroacetamide (BSTFA+TMCS, Sigma-Aldrich, St. Louis, MO) for 1 h at 80°C. The wax composition was analyzed by GC2010 gas chromatography (Shimadzu Inc., Kyoto, Japan) with the infector in splitless mode at the temperature programed from 80°C, 15°C per min to 200°C, 3°C per min to 300°C and 10 min hold at 300°C. Mass spectrum data were obtained on a GCMS-QP2010 mass spectrometer (Shimadzu, Kyoto, Japan) after impact ionization. The peaks were quantified using the GC-MS LabSolutions software (Shimadzu). Each wax monomer amount was determined based on internal standard and normalized by sampled seed surface area which was calculated from the total area of 1 mg seeds. Seed surface area was calculated as an ellipsoid surface area from the maximum and the minimum diameter of each seed determined by ImageJ software (Schneider et al. 2012).

### Polyester analysis

100 mg of mature seeds from simultaneously grown plants with three biological replicates were used for polyester analysis. Polyester extraction and analysis were performed by the method described previously with slight modification (Molina *et al*., 2006). Whole seeds were ground in liquid N_2_, delipidated and followed by methanolysis with sodium methoxide. Depolymerized compounds were silylated with BSTFA + TMCS (Sigma-Aldrich Inc.) for 20 min at 100°C and analyzed by GC-MS (GCMS-QP2010; Shimadzu Inc.) with the temperature programed from 140°C, 5°C per min to 310°C and 5 min hold at 310°C. The mass spectrum data were analyzed as mentioned above.

### Accelerated aging test

The accelerated aging test was conducted as previously described (Sattler *et al*., 2004). Seeds freshly harvested from individual four parental plants, as four biological replicates, were stored at 8 °C and 10% relative humidity for one week, and aged at 40 °C and 100% relative humidity for 72 h. Four sets of 60 seeds were used for each experimental plot. Each seed set was harvested from individual plants grown at the same time. Aged or untreated seeds were sterilized and their dormancy was broken using 5% Plant Preservative Mixture (Plant Cell Technology, Washington, DC) with 0.005% Tween-20 at 4 °C for 4 or 5 days. The seeds were germinated on Murashige and Skoog medium at 22 °C. The germination rate was calculated based on root emergence.

### Accession numbers

*LMI2* (AT3G61250), *MYB106/NOK* (AT3G01140), *MYB16* (AT5G13510), *FLP* (AT1G14350), *TfMYBML3* (AB649196), *AmMIXTA* (CAA55725), *AmMYBML1* (CAB43399), *AmMYBML2* (AAV70655), *AmMYBML3* (AAU13905), *PhMYB1* (CAA78386).

## Acknowledgments

We thank the Arabidopsis Biological Resource Center for providing the seeds for the T-DNA insertion lines. We also thank K. Tanaka (Saitama University) for technical support and advice regarding transmission electron microscopy, and T. Ishido, A. Kuwazawa, Y. Takiguchi, F. Tobe, A. Hosaka, K. Imokawa, K. Miyashita, K. Taguchi, Y. Sugimoto, and M. Yamada (AIST) for their technical assistance. We are grateful to K. Suzuki and Y. Naito (AIST) for advice related to the accelerated aging test. This work was partly supported by a Grant-in Aid for Young Scientists (B) (JP26840103) and a Grant-in-Aid for Scientific Research (C) (18K05447) from the Japan Society for the Promotion of Science and by JST, PREST Grant Number JPMJPR20D3 to Y. O.

## Author contributions

Y.O. conducted all experiments except for those in *T. fournieri* and some of the microscopic analyses. T.N. conducted the experiments in *T. fournieri*. Y.K. performed a part of fluorescent and electron microscopy experiments and analyzed the data. Y.O and T.I performed the lipid analysis. Y.O and N.M. designed all experiments and analyzed all data. Y.O. and N.M. wrote the article. M.K-Y. supervised the lipid analysis. N.M. and M.O-T. supervised the entire study.

## Short legends for Supporting Information

**Figure. S1** Meristem identity phenotype of 35S:LMI2-SRDX plants.

**Figure. S2** Phenotypes of 35S:LMI2-SRDX Arabidopsis thaliana and 35S:TfMYBML3-SRDX Torenia fournieri.

**Figure. S3** Phylogenetic tree of R2R3 MYB subgroup 9 proteins.

**Figure. S4** Phenotypes of immature seeds in the lmi2-3 mutnat.

**Figure. S5** Phenotypes of immature siliques and seeds in the progeny of the cross between wild type and lmi2-2 mutant.

**Figure. S6** Mucilage extrusion of mature seeds and cross section of fused seed in the lmi2-2 mutant.

**Figure. S7** Visualization of seed coat cuticle and cell wall of wild type and lmi2-2.

**Figure. S8** Seed wax monomer contents of wild type and lmi2-2.

**Figure. S9** Germination of wild-type, lmi2-2 and F2 seedsof them.

**Table S1** Frequency of observed phenotypes for 35S:LMI2-SRDX.

**Table S2** Branch number of Leaf trichome.

**Table S3** Primers used in this study.

