## Supporting Information for "LATE MERISTEM IDENTITY2 regulates cuticle formation on the seed surface and influences seed longevity"

Article acceptance date: Click here to enter a date.

The following Supporting Information is available for this article:

**Figure. S1** Meristem identity phenotype of *35S:LMI2-SRDX* plants.

**Figure. S2** Phenotypes of *35S:LMI2-SRDX Arabidopsis thaliana* and *35S:TfMYBML3-SRDX Torenia fournieri*.

**Figure. S3** Phylogenetic tree of R2R3 MYB subgroup 9 proteins.

**Figure. S4** Phenotypes of immature seeds in the *lmi2-3* mutnat.

**Figure. S5** Phenotypes of immature siliques and seeds in the progeny of the cross between wild type and *lmi2-2* mutant.

**Figure. S6** Mucilage extrusion of mature seeds and cross section of fused seed in the *lmi2-2* mutant.

**Figure. S7** Visualization of seed coat cuticle and cell wall of wild type and *lmi2-2*.

**Figure. S8** Seed wax monomer contents of wild type and *lmi2-2*.

**Figure. S9** Germination of wild-type, *lmi2-2* and F2 seedsof them.

**Table S1** Frequency of observed phenotypes for *35S:LMI2-SRDX.*

**Table S2** Branch number of Leaf trichome.

**Table S3** Primers used in this study.

**Fig. S1** Meristem identity phenotype of *35S:LMI2-SRDX* plants. (a) number of secondary inflorescences in *lmi2-2* mutant and *35S:LMI2-SRDX* and wild-type (Col.0) plants. Error bars represent standard error (n = 4 or 8). Single and double asterisks represent $p < 0.05$ and $p < 0.01$ by Student's t-test, respectively. (b) Phenotypes of *lmi2-2*, moderate and mild line of *35S:LMI2-SRDX*. Arrowheads indicate secondary inflorescences derived from the main stem. Bars indicate 5 cm.

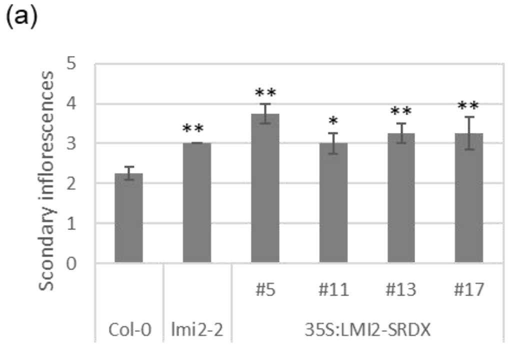


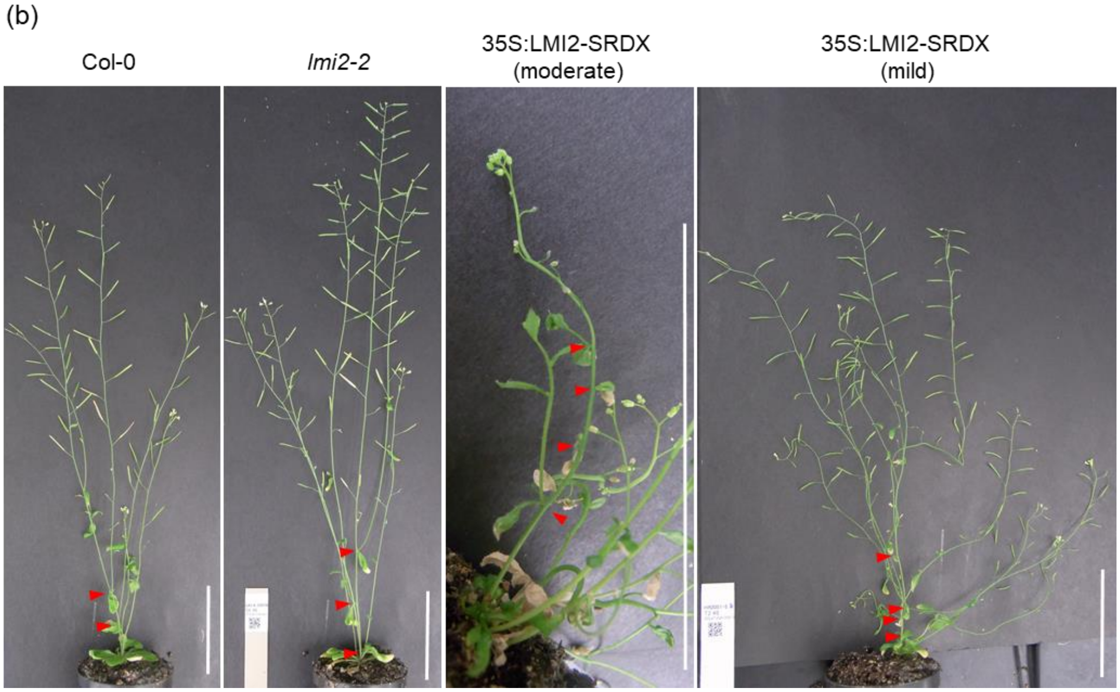

**Fig. S2** Phenotypes of 35S:LMI2-SRDX Arabidopsis thaliana and 35S:TfMYBML3-SRDX Torenia fournieri. (a and e) Vegetative (a) and reproductive (e) stages of wild-type Arabidopsis thaliana. (b) Fused leaves of a 35S:LMI2-SRDX A. thaliana severe-phenotype line. (c, d, g and h), Scanning electron micrograph of a flower (c and d) and stem (g and h) from wild-type (c and g) and 35S:LMI2-SRDX (e and h) plants. (f) Adhesion between a cauline leaf and a bud in a 35S:LMI2-SRDX A. thaliana mild-phenotype line. The arrow indicates the site of adhesion. (i and j), Wild-type A. thaliana (i) and 35S:LMI2-SRDX A. thaliana (j) stained with toluidine blue (TB). (k) Toluidine blue uptake per gram protein in A. thaliana. Error bars represent standard errors (n = 8). Double asterisks represent P < 0.01 according to Welch's t-test. (l and m) Wild-type T. fournieri (i) and 35S:TfMYBML3-SRDX T. fournieri (m) stained with TB. (n and o) Mature seeds of wild type (n) and LMI2pro:LMI2-SRDX(o). (p and q) Immature seeds of wild-type (p) and LMI2pro:LMI2-SRDX (q) plants stained with toluidine blue. Bars indicate 10 mm in (e) and (f), 5 mm in (a), (b), (i), (j), (l), (m), (n) and (o), 200 µm in (c) and (d), 100 µm in (p) and (q) and 10 µm in (g) and (h).

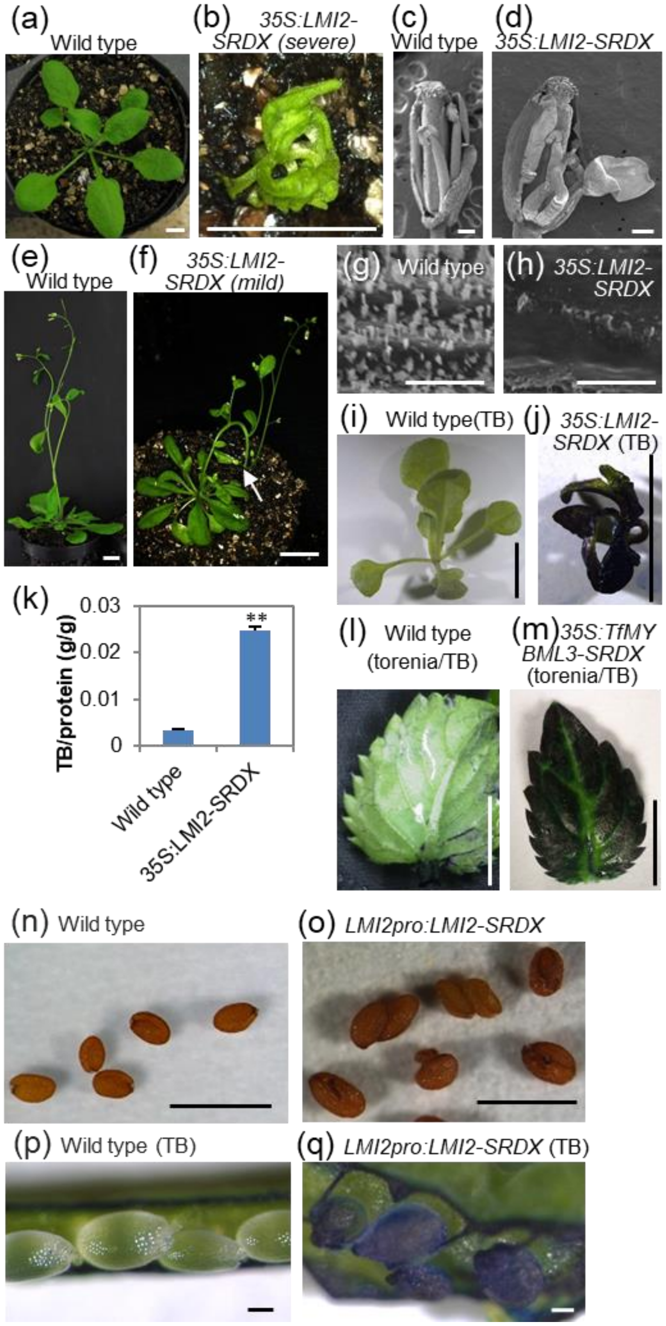
(a) Wild type
(b) 35S:LMI2-SRDX (severe)
(c) Wild type
(d) 35S:LMI2-SRDX
(e) Wild type
(f) 35S:LMI2-SRDX (mild)
(g) Wild type
(h) 35S:LMI2-SRDX
(i) Wild type(TB)
(j) 35S:LMI2-SRDX (TB)
(k)
TB/protein (g/g)
0.03
0.02
0.01
0
**
Wild type
35S:LMI2-SRDX
(l) Wild type (torenia/TB)
(m) 35S:TfMYBML3-SRDX (torenia/TB)
(n) Wild type
(o) LMI2pro:LMI2-SRDX
(p) Wild type (TB)
(q) LMI2pro:LMI2-SRDX (TB)

**Fig. S3** Phylogenetic tree of R2R3 MYB subgroup 9 proteins. The numbers next to the branches indicate percentage of replicate trees in which the associated taxa clustered together in the bootstrap test (1000 replicates). Bar with 0.1 represents evolutionary distance of 0.1 amino acid substitutions per site. “At” indicates *Arabidopsis thaliana*. “Am” indicates *Antirrhinum majus*. “Ph” indicates *Petunia x hybrid*.  “Tf” indicates *Torenia fournieri* Lind.

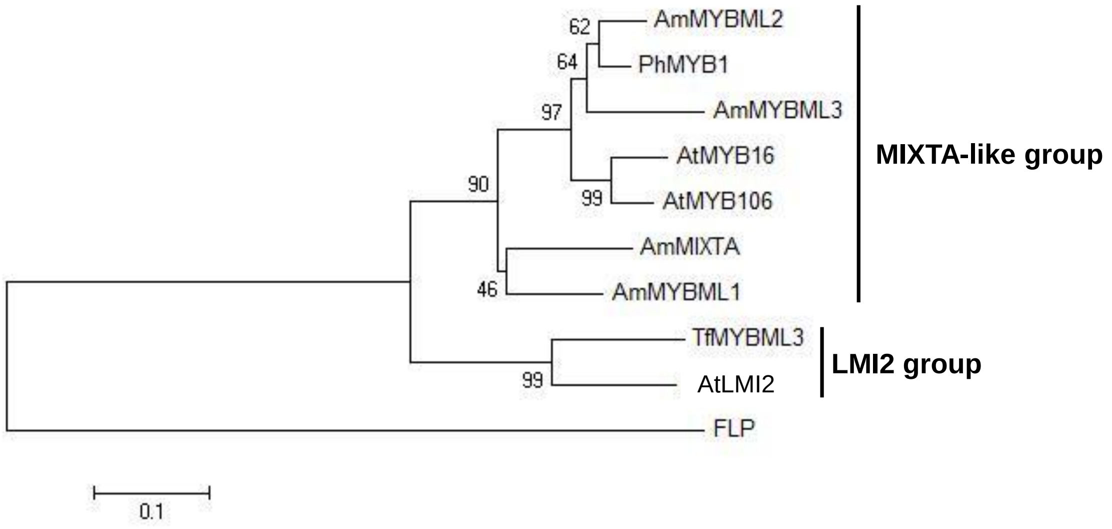

**Fig. S4** Phenotypes of immature seeds in the *lmi2-3* mutant. (a) Silique with average length of wild type and *lmi2-3*. (b, c) Inside of immature silique of *lmi2-3*. Red arrowheads indicate aborted seeds. Square brackets indicate seed clusters. Solid line arrow indicates torn funicle. (d) Average of silique length of all mature size silique on main stem in wild type and *lmi2-3*. Error bars represent standard error (n = 66 or 56). Asterisks represent p<0.01 by Welch's *t*-test. (e) Number of normal-size seeds, aborted and nonenlarged seeds in one immature silique of wild type and *lmi2-3*. Error bars represent standard error (n = 3). Seed number of each type is significantly different between wild type and *lmi2-3* (p<0.01 by Welch's *t*-test). (f) Fusion frequency in one silique of wild type and *lmi2-3*. Error bars represent standard error (n = 3). Asterisks represent p<0.01 by Welch's *t*-test. Bars indicate 5 mm in (a), 1 mm in (b), 100 µm in (c).

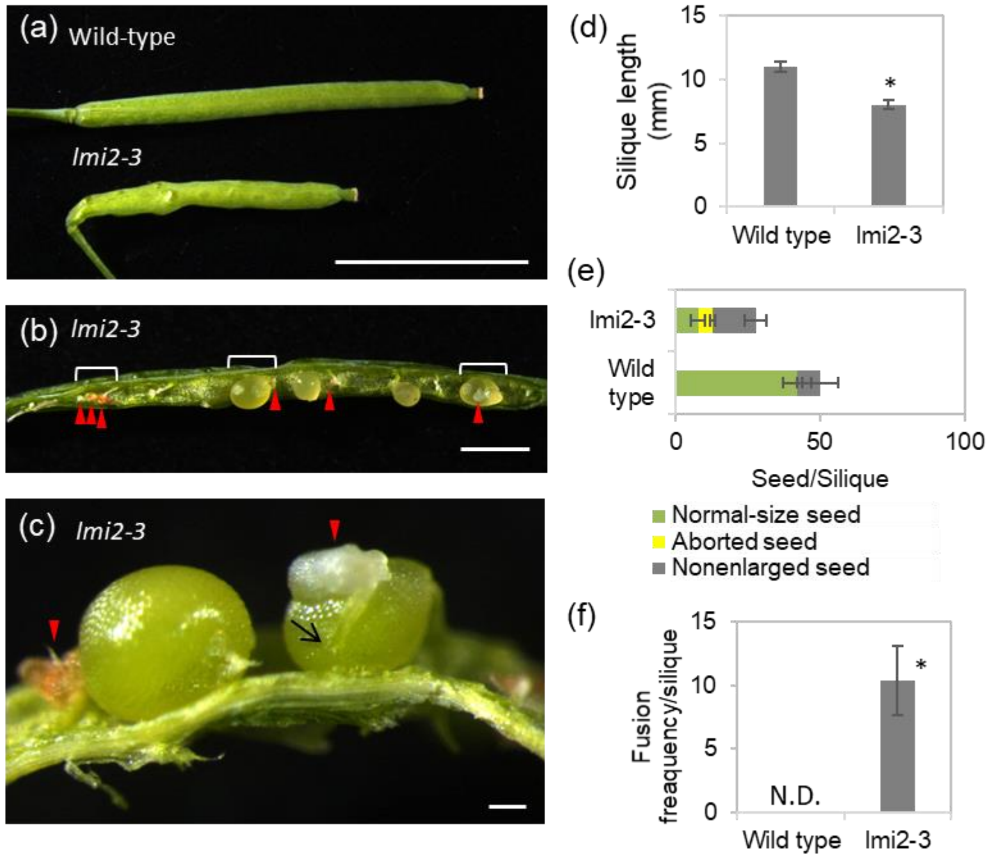

**Fig. S5** Phenotypes of immature siliques and seeds in the progeny of the cross between wild type and *lmi2-2* mutant. (a and b) Average length of all mature siliques on the main stem in $F_0$ (a) and $F_1$ (b) plants of indicated cross as “female (left) x male (right)”. Error bars represent standard error (n = 54 to 74). Asterisks represent $p<0.01$ by Welch’s *t*-test. (c) Representative silique with average length in $F_0$ and $F_1$ plants of indicated cross. (d and e) Average number of fused seeds in one silique from wild type and $F_0$ (d) and $F_1$ (e) plants of indicated cross. Error bars represent standard error (n = 3). Asterisks represent $p<0.01$ by Welch’s *t*-test. (f) $F_1$ and $F_2$ seeds in the immature silique of $F_0$ and $F_1$ plants of indicated cross. Arrowheads indicate fused seeds. (g) Confirmation of wild type (Col.0) and F1 plants. Genomic PCR was performed with *LMI2* and T-DNA specific primers. Bars indicate 5 mm in (c), 100 µm in (f).

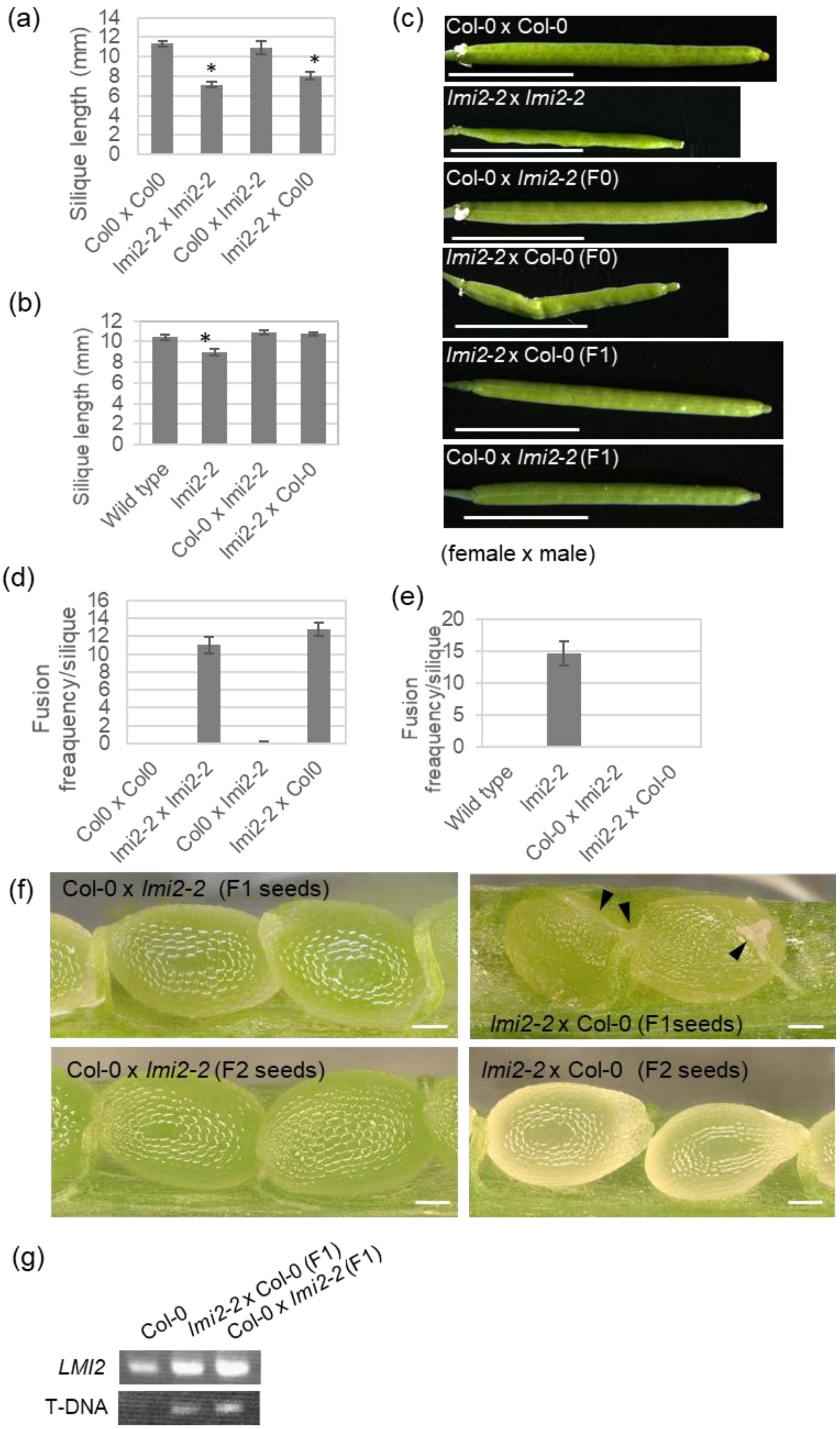
(a)
Silique length (mm)
Col0 x Col0
lmi2-2 x lmi2-2
Col0 x lmi2-2
lmi2-2 x Col0
(b)
Silique length (mm)
Wild type
lmi2-2
Col-0 x lmi2-2
lmi2-2 x Col-0
(c)
Col-0 x Col-0
lmi2-2 x lmi2-2
Col-0 x lmi2-2 (F0)
lmi2-2 x Col-0 (F0)
lmi2-2 x Col-0 (F1)
Col-0 x lmi2-2 (F1)
(female x male)
(d)
Fusion freaquency/silique
(e)
Fusion freaquency/silique
(f)
Col-0 x lmi2-2 (F1 seeds)
lmi2-2 x Col-0 (F1seeds)
Col-0 x lmi2-2 (F2 seeds)
lmi2-2 x Col-0 (F2 seeds)
(g)
Col-0
lmi2-2 x Col-0 (F1)
Col-0 x lmi2-2 (F1)
LMI2
T-DNA

**Fig. S6** Mucilage extrusion of mature seeds and cross section of fused seed in the *lmi2-2* mutant. (a-c) Seed coat mucilage of wild-type seeds (a) and *lmi2-2* seeds (b, c) were stained by ruthenium red. Asterisks indicate seeds stopped growing. (d, e) Cross section of wild-type (d) and *lmi2-2* (e) seeds stained with toluidine blue. Mucilage in epidermal cells were stained in blue. Some of them were extruded in aqueous solutions. Bars indicate 200 µm in (a-c) and 50 µm in (d, e).

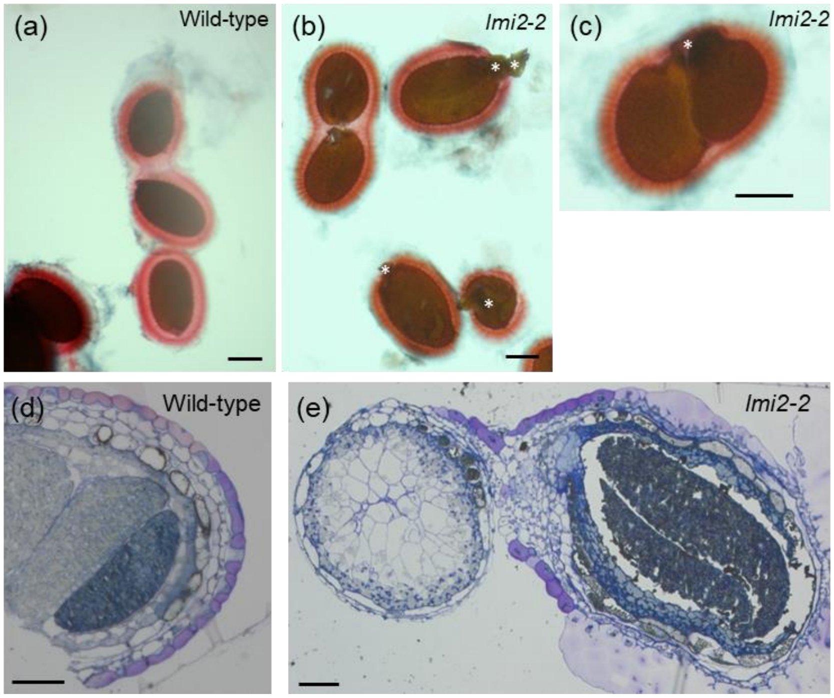

**Fig. S7** Visualization of seed coat cuticle and cell wall of wild type and *lmi2-2*. Cross sections of mature seeds from wild-type and *lmi2-2* plants were observed. Autofluorescence from aromatic compounds in cuticle, secondary cell wall, and the inside of seed (upper panel) and cell wall stained with fluorescent brightener 28 (middle panel) was observed under fluorescence microscope. Merged image is shown in bottom panel. White arrowhead indicates the fluorescence from cuticle outside cell wall in wild type seeds. *Lmi2-2* seed hardly had such fluorescence (black arrowhead). All pictures were shown in the same magnification.

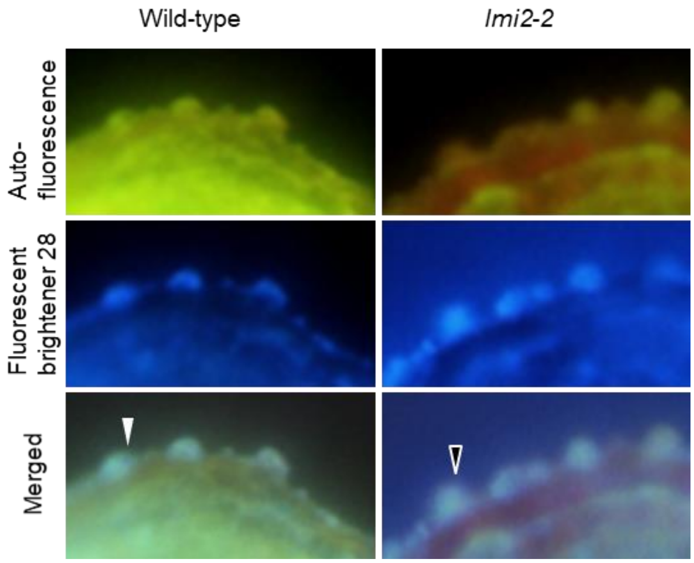

**Fig. S8** Seed wax monomer contents of wild type and *lmi2-2*. Alk; Alkane, PA; primary alcohol, SA; secondary alcohol, Ket; ketone, FA; fatty acid. Error bars represent standard error (n = 4 or 5). Single and double asterisks indicate $p < 0.05$ and $p < 0.01$ by Welch's *t*-test, respectively.

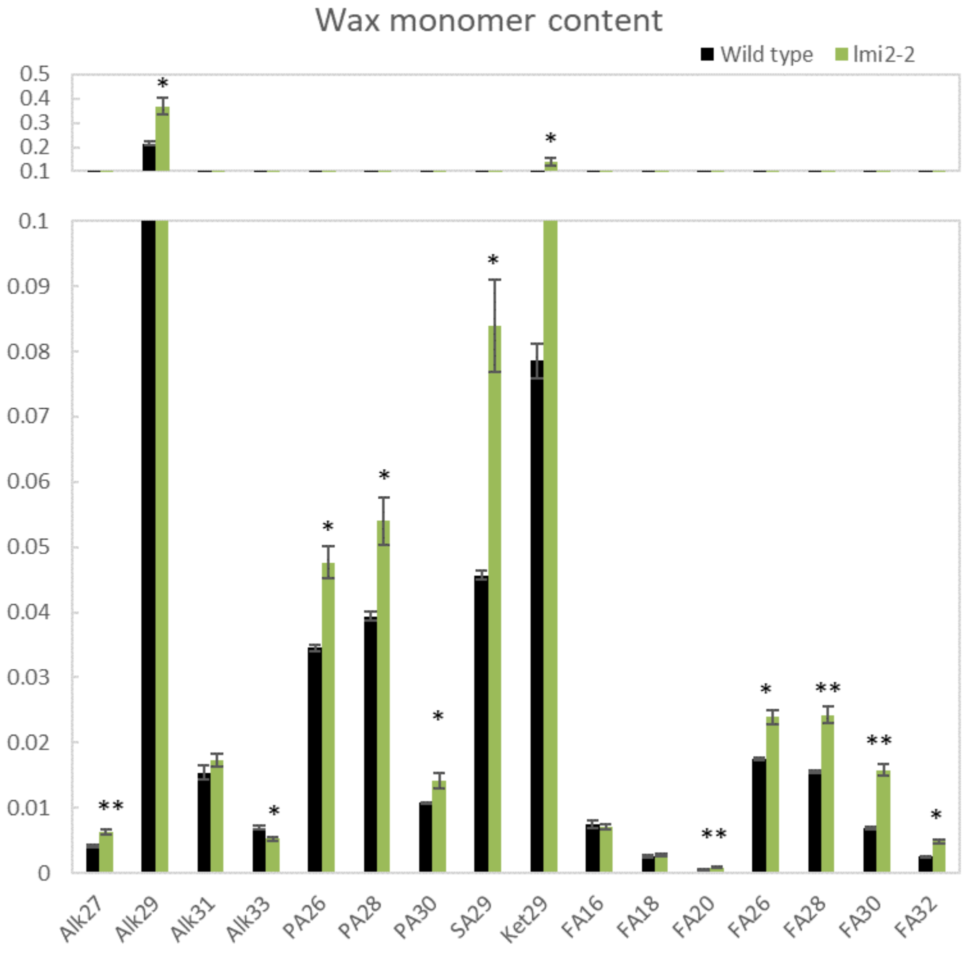

**Fig. S9** Germination of wild-type, *lmi2-2* and F2 seeds of them. Germination rate of untreated seeds (control) and seeds treated at 40 °C and 100% relative humidity for 3 days (accelerated aging). Error bars represent standard errors (n = 4). Single and double asterisks indicate P < 0.05 and P < 0.01 according to Welch's *t*-test, respectively.

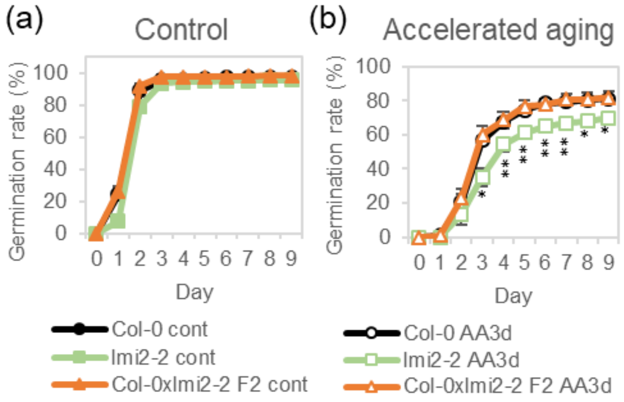


**Table S1.** Frequency of observed phenotypes for *35S:LMI2-SRDX.*

| | Severe | Mild | Wild-type like | Other | Total |
|---|---|---|---|---|---|
| Organ adhesion | 2 | 10 | 9 | - | 21 |
| Silique | 4 | 1 | 3 | 4 | 12 |

Severe, mild phenotypes about organ adhesion represent organ adhesions in whole plant, organ adhesion in limited tissues, respectively. Severe, mild and other phenotypes of silique represent low fertility and abnormal silique, abnormal silique, lethal before flowering, respectively. "Total" represents the number of examined plants for each transgene.

**Table S2.** Branch number of leaf trichome.

| | Branch number of trichome (%) | | | | | | Total* |
|---|---|---|---|---|---|---|---|
| | 1 | 2 | 3 | 4 | 5 | 6 | |
| Wild-type | 0.00 | 3.27 | 94.44 | 2.29 | 0.00 | 0.00 | 664 |
| *35S:LMI2-SRDX* | 3.50 | 36.00 | 60.00 | 0.50 | 0.00 | 0.00 | 200 |
| *lmi2-2* | 0.46 | 11.67 | 85.13 | 2.75 | 0.00 | 0.00 | 437 |

*Total number of trichomes counted on leaf 3 and 4 of 5 plants.

**Table S3. Primers used in this study.**

| Name | Sequence (5'-3') | Purpose |
|---|---|---|
| AT3G61250N | gATGGGAAGAACACCTTGTTGTGACAAGAT | *35S:LMI2-SRDX* |
| AT3G61250C | GAATTTGGAAACCATGGAAACAAGACCAAT | *35S:LMI2-SRDX, LMI2pro:LMI2-GFP:HSP* |
| AT3G61250pF3000 | GGGGACAACTTTGTATAGAAAAGTTGTCACGCTTATGGGTGTCACGCAATATT | *LMI2pro:GUS* |
| AT3G61250pR | GGGGACTGCTTTTTTGTACAAACTTGGCATTTGTTCCTCACCCCACTAACAAGC | *LMI2pro:GUS* |
| AT3G61250pFA | GGCGCGCCTCACGCTTATGGGTGTCACGCAATATT | *LMI2pro:GFP, LMI2pro:LMI2-GFP:HSP, LMI2pro:LMI2-SRDX* |
| AT3G61250pRB | CGCGGATCCTTGTTCCTCACCCCACTAACAAGC | *LMI2pro:GFP, LMI2pro:LMI2-SRDX* |
| AT3G10570pF1000 | GGGGACAACTTTGTATAGAAAAGTTGTTATCTTCCCGGAATTAGTGAAGACCC | *CYP77A6pro:LMI2-VP16* |
| AT3G10570pR | GGGGACTGCTTTTTTGTACAAACTTGGCATTTTTAGCTTCTTGTTTTTCTTCTT | *CYP77A6pro:LMI2-VP16* |
| 5G23940Rf1 | CAAAGACGCCGGAGTGAATTG | RT-PCR for *DCR* |
| 5G23940Rr1 | CCCGAAATCCACCTCGTAAACA | RT-PCR for *DCR* |
| 3G10570Rf1 | GAGCAAGAGCTCTCGAGGTT | RT-PCR for *CYP77A6* |
| 3G10570Rr1 | TCTCCGTCGTCTCTCGATGA | RT-PCR for *CYP77A6* |
| 1G51460.1Rf1 | TGGCGGATTTATGGCAGGCTTTA | RT-PCR for *ABCG13* |
| 1G51460.1Rr1 | TGCAACCCCATAGTGTCCATTGA | RT-PCR for *ABCG13* |
| 4G00360.1Rf1 | CGCCGGACGATGGAAAATTC | RT-PCR for *ATT1* |
| 4G00360.1Rr1 | GATCCTAGGTCCGGCGTTAA | RT-PCR for *ATT1* |
| 3G11430.1Rf1 | GCGGCTACGTTAGGGTTTGA | RT-PCR for *GPAT5* |
| 3G11430.1Rr1 | CGTTTCCAGCGAGAACCCTATA | RT-PCR for *GPAT5* |
| 1G53500.1Rf1 | TGGTTGAGGAGCTCTTGAGAGAAT | RT-PCR for *MUM4* |
| 1G53500.1Rr1 | TAGCGCGAGATCTTCGTGATGAA | RT-PCR for *MUM4* |

Underlines indicate attB sequences. Double under lines indicate restriction enzyme recognition site.
